# Genetic, morphometric, and molecular analyses of interspecies differences in head shape and hybrid developmental defects in the wasp genus *Nasonia*

**DOI:** 10.1101/663732

**Authors:** Lorna B Cohen, Rachel Edwards, Dyese Moody, Deanna Arsala, Jack H Werren, Jeremy A Lynch

## Abstract

Males in the parasitoid wasp genus *Nasonia* (*N. vitripennis, N. giraulti, N. longicornis*) have distinct, species specific, head shapes. Fertile hybrids among the species are readily produced in the lab allowing genetic analysis of the evolved differences. In addition, the obligate haploidy of males makes these wasps a uniquely powerful model for analyzing the role of complex gene interactions in development and evolution. Previous analyses have shown that complex gene interactions underpin different aspects of the shape differences, and developmental incompatibilities that are specific to the head in F2 haploid hybrid males are also governed by networks of gene interaction. Here we use the genetic tools available in *Nasonia* to extend our understanding of the gene interactions that affect development and morphogenesis in male heads. Using artificial diploid male hybrids, we show that alleles affecting head shape are codominant, leading to uniform, averaged hybrid F1 diploid male heads, while the alleles mediating developmental defects are recessive, and are not visible in the diploid hybrids. We also determine that divergence in time, rather than in morphological disparity is the primary driver of hybrid developmental defects. In addition, we show that doublesex is necessary for the male head shape differences, but is not the only important factor. Finally we demonstrate that we can dissect complex interspecies gene interaction networks using introgression in this system. These advances represent significant progress in the complex web of gene interactions that govern morphological development, and chart the connections between genomic and phenotypic variation.

## Introduction

Form develops in large part through the complex action and interaction of differentiating tissues and cells, and the gene regulatory networks (GRN) acting within them (Davidson *et al.* 2003). Stable changes in form within populations and between species are encoded by changes in the identity or magnitude of connections within and between developmental GRNs (Stathopoulos and Levine 2005; Hinman and Davidson 2007). Some interactions are relatively straightforward, resulting in phenotypes that are near to the expected sum or logical combination of the effects of the alleles alone in a neutral background. These are termed additive effects, and are often the result of independent pathways that contribute to a trait. In contrast, those phenotypes that are significantly different in magnitude and/or sign than the expected combination of alleles are due to the phenomenon of epistasis (Cheverud and Routman 1995). Epistasis among alleles is strongly indicative of direct interaction among the genes involved in producing the epistatic phenotype (Phillips 2008; Werren *et al.* 2016). Although some studies have argued that nearly all gene interactions are additive (Hill *et al.* 2008), a strong body of literature refute that claim, and even show that apparent additive effects can result from many underlying epistatic interactions (Avery and Wasserman 1992; Cheverud and Routman 1995; Huang *et al.* 2012; Jones *et al.* 2014). This discrepancy is likely due to detection bias, whereby statistical constraints limit testing to pairwise interactions, or search for quantitative trait loci (QTL) only among regions that show a significant marginal effect (Wolf *et al.* 2000). In fact, much epistasis involves chromosome regions that show little marginal effects, and three- or four-way interactions are quite common (Templeton *et al.* 1976). Non-biased epistatic QTL methods face much greater technical challenges than individual QTL mapping methods, but can be very informative as they simultaneously weigh the mean additive or non-additive effects on phenotype (Carlborg and Haley 2004).

While these nonlinear genetic interactions complicate the genotype-to-phenotype map, they are essential in generating complex and quantitative traits. Knowledge of epistatic interactions will deepen our understanding of complex traits, how they are encoded in the genome, and how they evolve (Mackay 2014). Thus investigating developmental GRNs is crucial to understand the genetic basis of form (Phillips 2008). The head is perhaps the most complex morphological feature of the bilaterian body plan. Considerable developmental challenges in patterning and development are encountered as several major sensory organs arise from a common primordium, like the eye-antennal disc, (Domínguez and Casares 2005; Palliyil *et al.* 2018), and then integrate with other primordia, such the labial disc and gnathal segments (Younossi-Hartenstein *et al.* 1993).

This complexity is reflected in the gene networks underlying head development, and the complex genetic interactions that participate in head development revealed by the highly epistatic nature of pathological syndromes in humans and model organisms (Lidral and Moreno 2005; Wolf *et al.* 2005; Gross *et al.* 2014). Since these model systems are standard diploids, analysis of complex epistatic interactions suffers from the complications of dominance effects and extremely rapid increase in the number of progeny required to detect gene interactions (Werren *et al.* 2016). Epistatic interactions among multiple recessive alleles are quite demanding to detect due to the exponentially increasing rarity of progeny homozygous for the required alleles at all loci involved. In diploid organisms the rate of obtaining the correct genotype is ¼^X^ for recessive interacting alleles, where x is the number of loci involved in the producing the epistatic phenotype (Werren *et al.* 2016).

Conversely, use of a haploid model system significantly increases the frequency of the (1/2^x^ for haploids vs 1/4^x^ for diploids) and eliminates interference from dominance effects. The preceding is a major reason why haplodiploid insects in the genus *Nasonia* show strong promise as model systems for understanding how epistasis and complex interactions among alleles in GRNs that affect the evolution of form (Gadau *et al.* 2002; Hoedjes *et al.* 2014). Like all Hymenoptera, *Nasonia* have haplodiploid genetics, where unfertilized eggs obligate haploid males, and fertilized eggs become diploid females (Werren and Loehlin 2009). Additionally, *Nasonia* are small insects easily reared in the lab, have short generation times, can be kept alive under refrigeration for extended periods, have fully annotated genomes, visible and molecular markers and crucially, the ability to make viable, fertile F1 hybrids between all species (Werren and Loehlin 2009; Lynch 2015). Furthermore, there are clearly marked morphological differences between the species, particularly among the haploid males, making evolutionary genetic analysis possible in *Nasonia* (Beukeboom and Desplan 2003; Werren and Loehlin 2009; Lynch 2015; Werren *et al.* 2016).

The distinctness of male head morphology is particularly apparent in the males of *N. giraulti* (Werren *et al.* 2016), (Figure 1, Figure 2A-E, Table S1). Females of all species of the genus, and males of *N. vitripennis* have a rounded ovoid face shape. In contrast, *N. giraulti* male faces are mostly square, with consistent width along the length of the face (Figure 1, TableS1). The exception to this square-ness is the cheeks, which protrude ventrally, giving these males a jowly appearance (Werren *et al.* 2016), (Figure 1, Figure 2E, Table S1).

**Figure 1.**
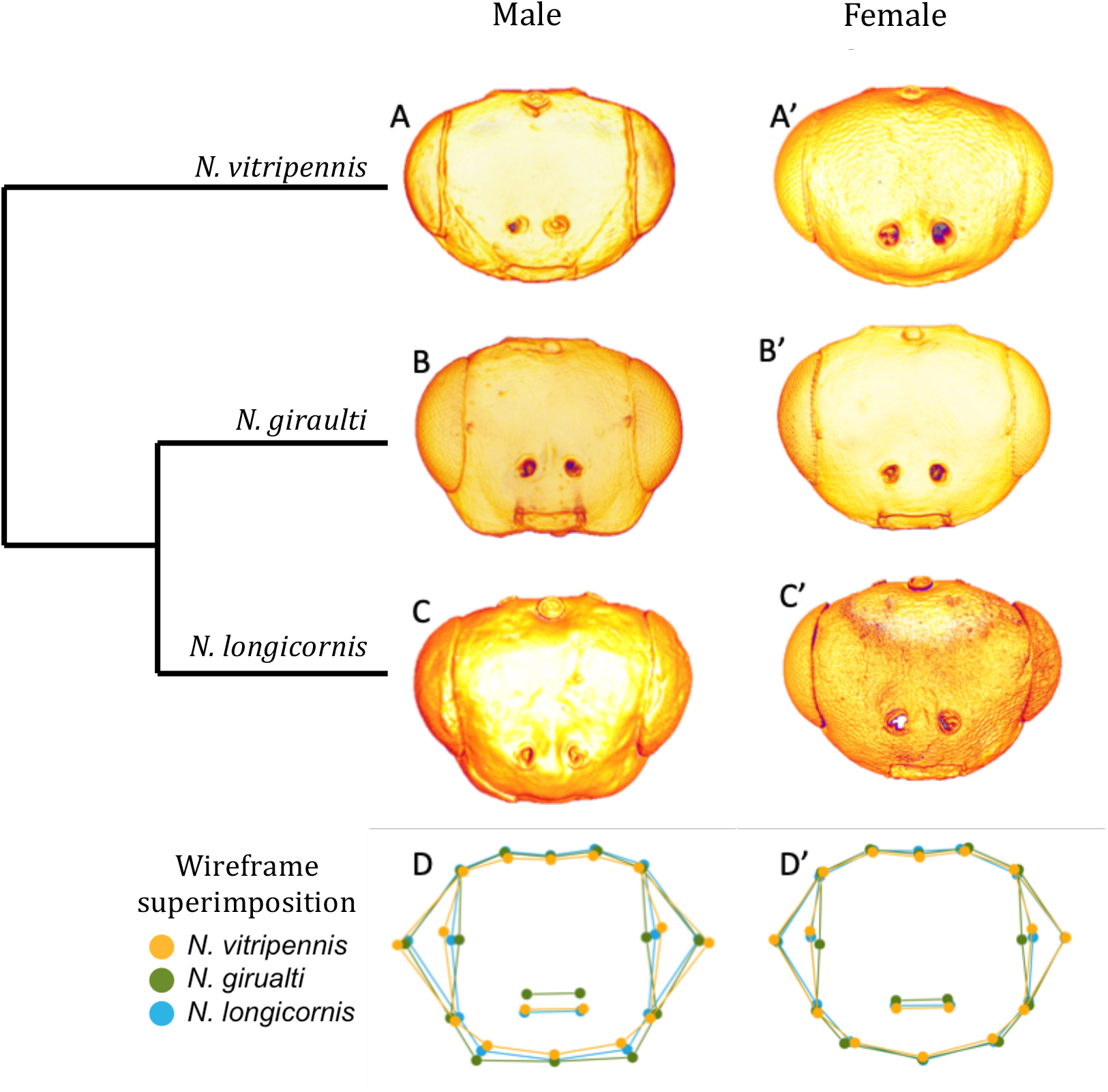
Shape differences among wild type species. A-C’) Representative images of wasp heads. D-D’) Procrustes superimposition of average wild type head shapes based on 16 landmarks. Morphology recapitulated by wireframe diagram. A) *N. vitripennis* male, A’) *N. vitripennis* female, B) *N. giraulti* male, B’) *N. giraulti* female, C) *N. longicornis* male, C’) *N. longicornis* female, D) Superimposed wireframe diagrams of male heads D’) Superimposed wireframe diagrams of female heads. Yellow landmarks denote *N. vitripennis*, green *N. giraulti*,and blue *N. longicornis.*

**Figure 2.**
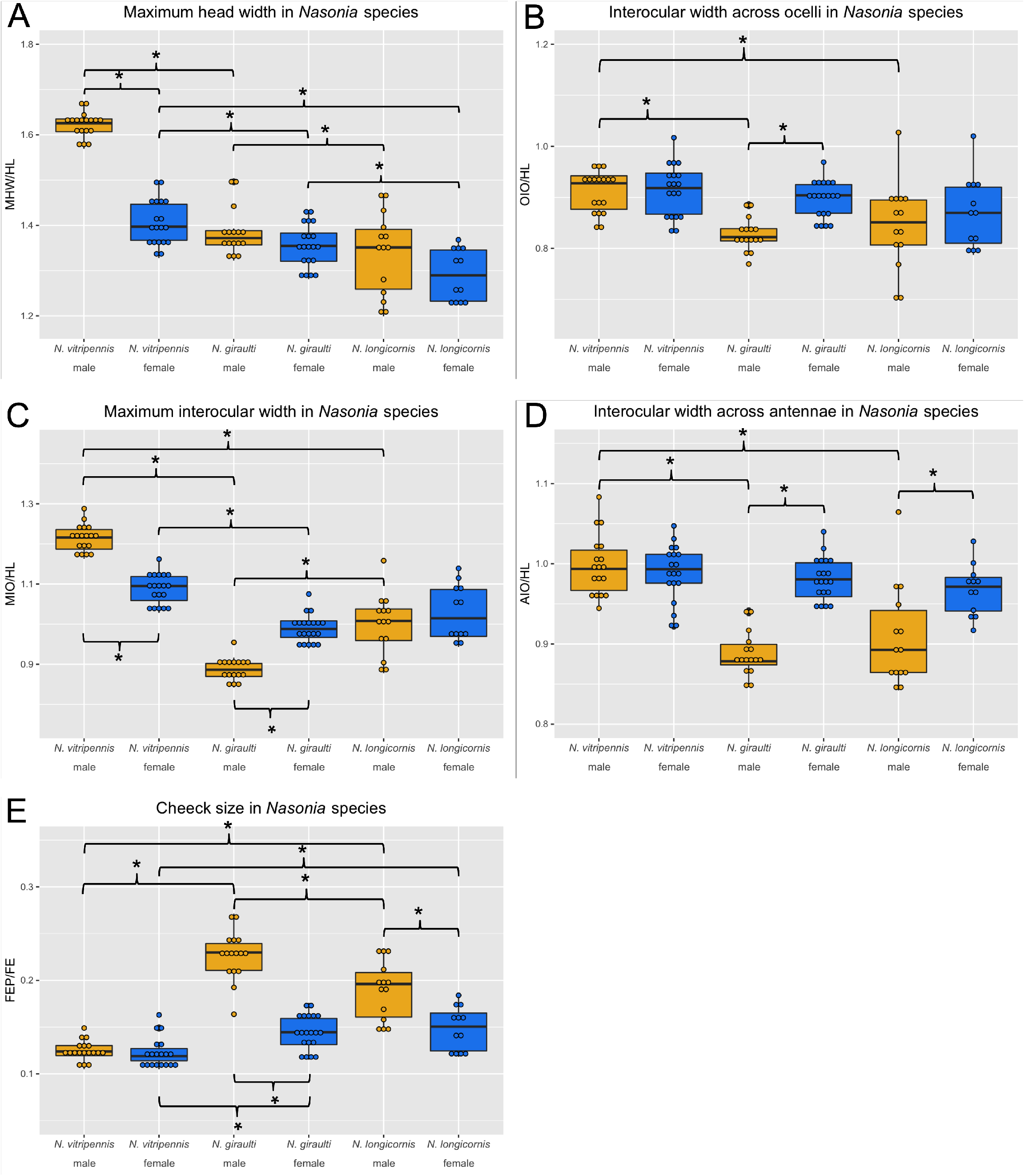
Measurement ratios of each parent species presented as box and whisker plots. Each dot represents a single individual, a box represents the inter-quartile range, the center line represents the median value and vertical lines represent upper and lower quartile ranges. A) Maximum head width over head length (MHW/HL), B) Interocular width at ocelli over head length (OIO/HL), C) Maximum interocular widther over head length (MIO/HL), D) Interocular width at antennae over head length (AIO/HL), E) Cheek size (FEP/FE.) Males are shown in yellow and females in blue. Comparisons were made among males of each species, among females of each species, and between males and females within each species. Asterisks indicate P<0.05.

In addition to functional hybrid males with mixtures of morphological features found in the males of the parental species, F2 male hybrid offspring between *N. vitripennis* and *N. giraulti* display a wide variety abnormal phenotypes (Werren *et al.* 2016). These defects include cranial midline furrowing, dorsal-ventral asymmetries, and lateral asymmetries (Werren *et al.* 2016). Preliminary QTL analyses indicate that all of these abnormalities are largely due to epistatic interactions among alleles of several genes from the two species (Werren *et al.* 2016).

Here, we aim to develop a better understanding of the genetic and developmental origin of the phenotypes we observe between species and among hybrids. Our analyses address several outstanding questions about the nature of the head patterning GRNs of the two species and how alleles interact to produce different hybrid phenotypes by combining interspecies crosses, RNA interference, and cross-species introgression analyses. Questions we address include: 1) Are development defects in hybrid F2 males correlated with divergence time between the species or with degree of morphological divergence? 2) Are the defects due to general developmental instability or to disruption of gene interactions specific to the head? 3) Are the defects primarily due to the exposure of allelic incompatibilities in haploids, or are there dominant alleles involved in the formation of novel structures and shapes arising between the species? We also address the dominance relationships among alleles affecting head shape and the developmental defects observed in hybrid males.

## Materials and Methods

### Hybrid crosses

Wolbachia-free and highly inbred strains of *N. vitripennis* (AsymCx), *N. giraulti*, (RV2x) and *N. longicornis* (IV7) (Werren *et al.* 2010) were used to make hybrids. For each cross a ratio of fifteen females to nine males were allowed 24 hours to mate before provided fly hosts to parasitize. Fifteen to twenty F1 hybrid virgin females from each interspecies cross were then provided hosts to parasitize. Setting females as virgins guarantees all offspring to be haploid males.

### Measurements

Heads from male hybrids were stained, mounted and imaged as outlined by Werren *et al.* 2016. Measurements were taken in Imaris7.1.1 according to parameters also outlined in Werren *et al.* 2016. Acronyms are as follows: MHW-maximum head width, HL-head length, OIO-interocular distance through ocelli, MIO-maximum interocular distance, AIO-interocular distance across antennal sockets, FE-distance from bottom of eye to center of mandible, FEP-farthest point on cheek perpendicular to line FE (see Figure S1 for diagram). Measurements are presented as ratios to normalize natural difference in overall size of the individual. MHW, OIO, MIO and AIO are normalized in relation to head length (HL) and dividing FEP by FE normalizes cheek size. Mann-Whitney U tests were performed for nonparametric data between two groups, and Bonferroni adjustments made for multiple comparisons. For wild type parent species, comparisons were made among males of each species, among females of each species, and between males and females within each species. Each experimental group was compared individually to *N. vitripennis* and *N. giraulti* wild type males. Plots were generated using R (R Core Team 2013), raw averages, standard deviations and significance can be found in Tables S1 and S3.

### Analyses of symmetry

#### Heads

Head symmetry was measured by Procrustes distance analysis of 105 hybrid male heads as well as 72 wild types (30 *N. vitripennis* and 28 *N. giraulti* of equal males and females). Each head was marked at 16 landmarks: One at each ocellus, at the tops and bottoms of each eye, at the maximum arc of each eye, the maximum arc of each cheek, the center point of the mandible, both ends of the MIO, and location of each antennal socket (Figure S1). Landmarks were established three times for each head and coordinates for each landmark were averaged and imported as an array in R (R Core Team 2013). Scaling, rotating and superimposition of head landmarks was carried out using R packages gemorph, shapes and Momocs (Bonhomme *et al.* 2014; Adams *et al.* 2017; Dryden 2017). R package vegan (Oksanen *et al.* 2017) quantifies symmetry by overlaying the left and right sides of heads and performs Procrustes distance analyses, defined as **Σ**((distance between corresponding landmarks)^2^).

#### Legs and wings

Front wings and T1 legs of the same 105 hybrid and 72 wild type wasps were carefully removed and mounted on slides. Each wing and leg was imaged on a Zeiss Stereo Discovery V.8 dissecting scope using Zeiss Axiovision software v. 4.8. Each specimen was measured three times and the length averaged. The difference in length between left and right sides of each appendage was compared for hybrids and wild types.

### RNAi

#### Diploid males

To generate diploid males, we used parental RNAi (Lynch and Desplan 2006) on a mutant strain of *N. vitripennis* with grey eye color (*N.vit^Oy/Oy^*). Female yellow pupae of *N.vit^Oy/Oy^* were injected with 1ug/ul of double-stranded RNA (dsRNA) targeting *Nv-transformer (Nv-tra)*, whose function is required for female development in fertilized eggs (Verhulst *et al.* 2010). The injected *N.vit^Oy/Oy^* adult females were then crossed to the wild type *N. giraulti* (RV2x), which have a red-brown eye color. While haploid males display the grey eye phenotype, the hybrid, diploid males express wild type red-brown eye color allele obtained from the *N. giraulti* parent. Male vs female offspring are easily differentiated in the pupal state by wing size and absence/presence of an ovipositor (Werren and Loehlin 2009).

Primer Sequences (Arsala and Lynch 2017):

*Nv*-Transformer-F: ggccgcgggcaaaatccgtgagacaac

Nv-Transformer-R: cccggggcgaggctgtcggcaaaaata

#### Dsx

Knockdown of *N. giraulti doublesex (Ng-dsx)* was carried out by injecting *N. giraulti* larvae with dsRNA targeted to *Ng-dsx* according to (Werren *et al.* 2009). Mid-stage larvae collected ~8 days after egg lay were positioned on double-sided tape on a slide for injection. Larvae were returned to 25° incubator to eclosion. Adult heads were stained, imaged and measured as described above.

*Ng*-Doublesex-F: ggccgcggcgcggaaagttgaagaagtc

*Ng*-Doublesex-R: cccggggcaatccaagtcccacatctgc

### Introgression

Introgression of *Ng* chromosomal regions into an *Nv* genetic background is routinely used to investigate the genetic basis of differences in traits between the species, and some cases for positional cloning of causal loci. We generated an *Ng* introgression into *Nv* of a region on chromosome 2 implicated in abnormal head clefting in F2 males (Werren *et al.* 2016). The initial chromosome 2 introgression line is designated INT_2C, and head shape effects were observed, in addition to phenotypic effects on body color, survival and female fertility (data not shown). Subsequent recombinants where generated by using primers flanking insertion/deletion differences across the region. A smaller scale introgression designated 2C-Cli was produced that shows an abnormal head clefting in both males and females. The recessive lethal and female fertility effects were separated from the clefting region by recombination. Both introgression lines were generated according to methods described in Breeuwer and Werren, 1995. The smaller region is estimated to be 16 centimorgan based on the map in Desjardins *et al.* 2013. The line with an introgression on chromosome four (denoted INT_wm114) was generated to study the sex-specific gene size differences in *Nasonia* (Loehlin *et al.* 2010). Adult heads were stained, imaged and measured as described above.

## Data Accessibility

Strains are available upon request. Figure S1 shows how heads were measured. Table S1 provides the raw measurements of the parental species heads. Table S2 gives the measurements of the wings and legs of parental species and hybrid wasps. Table S3 provides the measurements of the experimental strain heads. Table S4 provides a side by side comparison of the measurements of parental and experimental heads. The authors affirm that all data necessary for confirming the conclusions of the article are present within the article, figures, and tables.

## Results

### Wild type males have species-specific morphologies

The significant differences in head shape between the males of *N. vitripennis* and *N. giraulti* were described in Werren *et al.* 2016, and a general description of *N. vitripennis, N. giraulti*, and *N. longicornis* heads was provided in Darling and Werren, 1990. To understand how head shape has evolved in the *Nasonia* genus, we examined head shape of the males and females in more detail. To this end, we took seven measurements (Figure S1) on several heads (n=12-20, Table S1) from both males and females of the three species we investigate here. These measurements include maximal head width, head length, facial width in three places, and cheek size.

General head shape measurements were normalized by dividing the measurement by the head length to consistently control for variation in head size. Cheek size measurements are expressed as ratios with another measurement of head size to normalize for natural size variation across individuals and since females tend to be larger than males. Upon comparing measurement ratios, we found that *N. vitripennis, N. giraulti* and *N. longicornis* females all have roughly the same oval shape in their faces (Figure 1A’, B’, C’, D’, Figure 2B-D), with *N. giraulti* females having the least round shape (Figure 2C). Males on the other hand, all look quite different (Figure 1A, B, C, D). The males of *N. vitripennis* look nearly the same as that of the females (Figure 1A-A’), but are actually wider at maximum head width (MHW) and maximum interocular distance (MIO) (Figure 2A and C). The males of *N. giraulti* (Figure 1B) appear much more square due to their consistency in the three normalized face width measurements: across ocelli (OIO), across antennae (AIO), and MIO (Figure 2B-D, Table S1). Male heads of *N. longicornis* (Figure 1C), were previously described to be similar to *N. vitripennis* males and females (Darling and Werren 1990), but actually measure more similar to *N. giraulti* in terms of face shape at measurements MHW, OIO and AIO (Figure 2A, B, D, Table S1). Additionally, *N. longicornis* males are almost exactly intermediate between the other two species in cheek size (FEP/FE) (Figure 1D, Figure 2E, Table S1).

Also interesting to note is that a few traits are partially sex specific. For example, females of *N. giraulti* and *N. longicornis* do have bigger cheeks than *N. vitripennis*, but the males traits are extreme (Figure 2E). This implies incomplete sex specificity of the shape differences between species, and that some genes responsible for the extreme male differences also affect female development. Similar phenotypes appear to exist at the interocular width at the top and bottom of the head, OIO and AIO, where both male and female *N. giraulti* and *N. longicornis* have relatively narrower faces than the *N. vitripennis* counterparts, but again the male trait difference are more exaggerated. Overall conclusions from wild type species shape analyses is that females across the three species have roughly the same shape except for maximum head width, intraspecies sex-specific differences are starkest within *N. giraulti*, and that *N. longicornis* displays phenotypes intermediate between the other two species, in contrast to what was previously reported (Darling and Werren 1990; Werren *et al.* 2010).

### Developmental incompatibility alleles more strongly associated with temporal, rather than morphological divergence

Understanding the genetic basis of morphological divergences can provide insight to how morphology evolves. One question that needs to be addressed is whether the genetic basis of developmental defects in F2 hybrids between *Nv* and *Ng* are caused by negative interactions among alleles involved in producing the divergent head shapes. One could imagine that alleles important for making the exaggerated traits may contribute to tissue behavior incompatible with the effects of alleles driving the formation of elongate, ovoid head of *Nv*. Similarly, alleles required to produce the novel *Ng* cheeks may have unexpected interactions with alleles of the cheek-less *Nv*. On the other hand, the F2 hybrid male head defects could occur due to interactions among divergent alleles that have changed due to forces other than morphological evolution of the head, such as random drift over the course of the 1.4-1.6 million years of independent evolution since these species shared a common ancestor.

To differentiate between these possibilities, we examined F2 hybrid males created with *N. longicornis* (*Nl*). *Nl* is a sister species to *N. giraulti*, from which it diverged ~0.4 - 0.56 million years ago (mya). The divergence time between *Nl* and *Nv* is identical to that between *Ng* and *Nv* (~1.4 million years). We have found that, while male *Nl* heads are significantly less square, and have significantly smaller cheeks (Figures 1 and 2, Table S1) than *Ng* males, they are also statistically significantly different from *Nv* in these measures (Figures 1 and 2, Table S1). Thus, divergence time is not completely uncoupled from morphological evolution in this experiment. However, the timing of the origin of the negative interactions leading to developmental defects can still be inferred as could a potential influence of the exaggerated morphological differences in *Ng* relative to *Nv*.

Eighty-eight percent of F2 males produced by *N. giraulti* x *N. vitripennis (Ng-Nv)* hybrid females show head defects of some type, while 80% of F2 hybrid males resulting from *N. longicornis* x *N. vitripennis* hybrids (*Nl-Nv*) exhibit head defects (Figure 3A). These defects took many forms, with some co-occurring in the same individual. Lateral asymmetry indicates a difference in relative size between the left and right sides of the head (Figure 3B, compare arrows), or misplacement of ocelli (Figure 4A). Individuals with a cleft phenotype display a furrow among the midline of the face (Figure 3B’ arrowhead). In wild type wasps, the point at which the eye meets the epidermis at the top of the head is directly above the point where the eye meets the epidermis at the bottom of the head (Figure 1D-D’). When this is not the case in hybrid individuals it is referred to as dorso-ventral (DV) asymmetry (Figure 3B’, compare arrows), which can occur in one or both eyes. Abnormalities that account for less than five percent of the hybrid population are grouped under “miscellaneous.” These include swollen head syndrome, an expansion at the top of the head (Figure 3B’’); bulging eye syndrome, where the eye field is larger than average causing the facial area to be smaller than average; pitting around the antennal sockets; and presence of a fourth ocellus. Some individuals display more than one type of abnormality, which are noted under “multi” in Figure 3A. In contrast to the high rate and diversity of head defects in the *Nl-Nv* and *Ng-Nv* F2 male hybrids, observable head defects are seen in only ~20% of the F2 *N. longicornis* x *N. giraulti* (*Nl-Ng*) hybrids (Figure 3A). Strikingly, the clefting phenotype was completely absent and both DV and lateral asymmetries only occurred in five percent of hybrids (compared to ~25% and 20% in hybrids involving *Nv*, respectively, Table 1). Miscellaneous defects accounted for 10% of abnormal heads in *Nl-Ng* F2 hybrid males (compared to 18-24% in *Nv* hybrids, Table 1) and no individuals of this cross had more than one defect (compared to 10-12%of *Nv* hybrids, Table 1).

**Figure 3.**
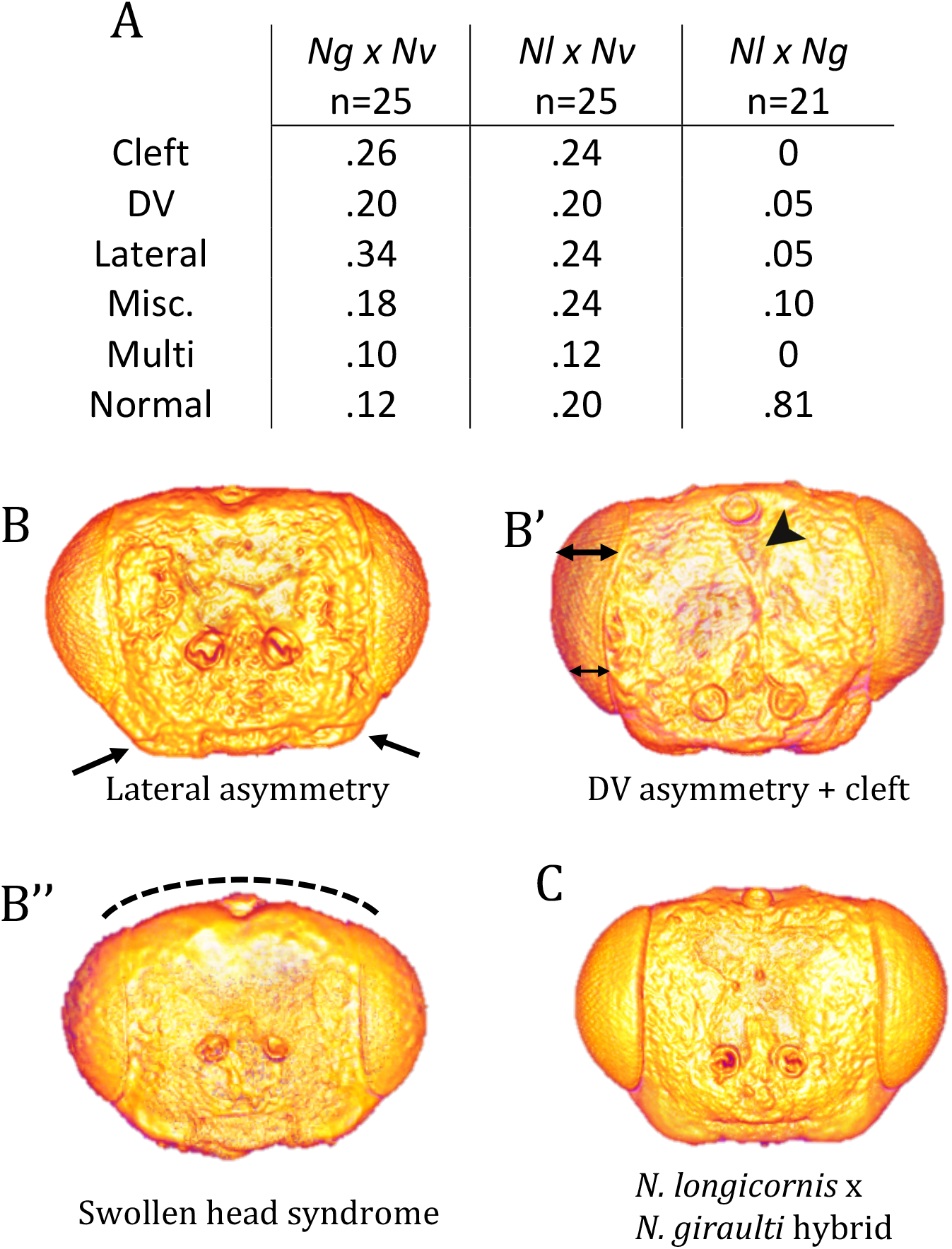
Representative hybrid head shapes from N. longicornis crosses. A) Table containing percentages of hybrid offspring that display each category of facial defect for the three hybrid crosses. The first three categories are facial clefting, dorsoventral asymmetry, and lateral asymmetry. Individuals displaying more than one type of defect are noted under Multi. Miscellaneous defects include swollen head syndrome, bulging eye syndrome, and antennal pits. B-B’’) *N. longicornis* x *N. vitripennis* hybrids. B) Lateral asymmetry, arrows points to difference in cheek size. B’) DV asymmetry and midline cleft, double-ended arrows indicate chance in width of eye field from dorsal to ventral side of the head. Arrowhead points to midline cleft. B’’) Swollen head syndrome, the top of the head bulges outward. C) *N. longicornis* x *N. giraulti* hybrid. Note no obvious aberrations.

In summary, it appears that most of the alleles causing developmental defects in the heads of hybrids between *N. vitripennis* and *N. longicornis* or *N. giraulti* arose and were fixed prior to the divergence of the *N. giraulti* and *N. longicornis* lineages from each other ~ 400k-500k years ago (Campbell *et al.* 1993; Martinson *et al.* 2017). This indicates that exaggeration of morphological differences in *N. giraulti* males had little effect on the evolution of developmental incompatibility between *N. giraulti* and *N. vitripennis*. The low frequency of defects seen in *Nl-Ng* hybrids may be due to new alleles that have arisen in one or both lineages, or may reflect independent sorting of polymorphisms present in the ancestral population that gave rise to them.

### Asymmetric hybrid phenotypes are specific to the head

The most common of the abnormal hybrid phenotypes is asymmetry (Figure 3A). We wanted to know the extent of these asymmetries and whether they are caused by a general developmental instability in the hybrids, as is often seen in some systems (Alibert and Auffray 2003; Leamy and Klingenberg 2005), or if the phenotype has its basis in genetic mechanisms operating specifically in the head. To determine this, we developed an approach to quantify asymmetry among head capsules as well as difference in length at two other body parts: legs and wings (Figure 4E). Symmetry between left and right sides of heads was quantified using R package vegan (Oksanen *et al.* 2017), wherein landmarks from the left are overlayed to their corresponding landmarks on the right (ie, the wireframe is folded along the centerine) and a Procrustes distance analyses is performed by calculating **Σ**((distance between corresponding landmarks)^2^). A Procrustes distance analysis (Figure 4A-C) done on 105 *Nv* x *Ng* hybrid heads found that a hybrid head has only 93% correlation on average between its left and right sides (Figure 4D). On the other hand, wild type heads measured from both males and females of *N. vitripennis* and *N. giraulti* revealed a 99.5% correlation between left and right sides of the head. The differences in correlation are highly statistically significant (P<0.001), indicating a strong effect of the hybrid genome on the maintenance of tissue homeostasis. However, we found no significant difference in the length between the left and right T1 legs, nor between the first pair of wings in the same set of F2 hybrid wasps, as compared to either parental species (Figure 4E, Table S2). We therefore conclude that generalized developmental instability is not a likely explanation for cranial asymmetry, since we do not observe defects or asymmetries in other body parts. Rather, there appears to be phenomenon specific to the head patterning and homeostasis system.

**Figure 4.**
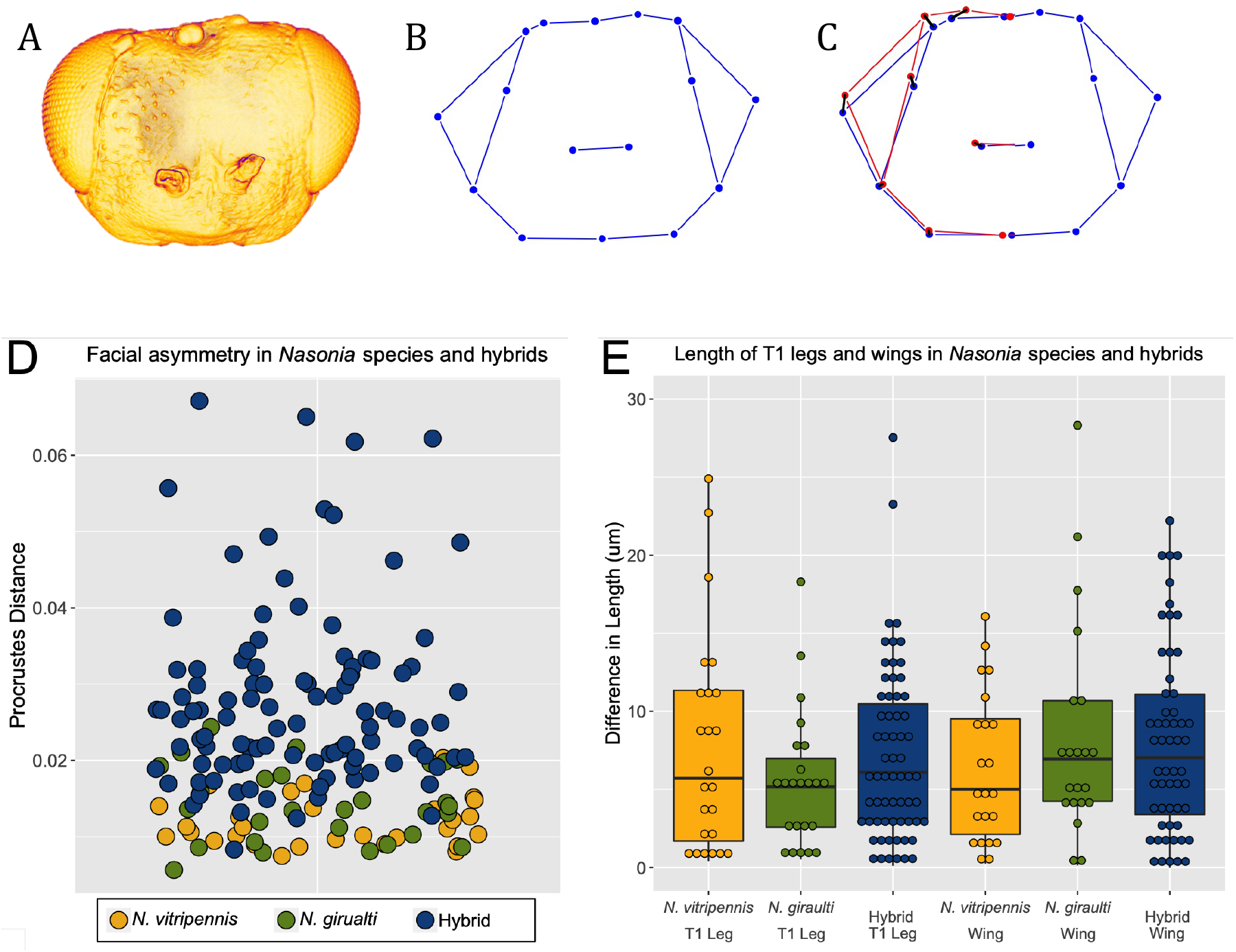
Symmetry analyses. A) Representative asymmetric hybrid head. B) Wireframe diagram of head in (A). C) Right-side landmarks reflected over left side landmarks. Reflection is shown in red. A black line represents distance between corresponding landmarks. Procrustes distance is calculated as the sum of the squares of each distance. D) Scatter plot in which each dot depicts Procrustes distance for individual wasps. Dark blue dots represent hybrid individuals; yellow, green and light blue are wild types. P<0.001 between hybrids and wild types. E) Box Plot graphing differences in length of T1 legs and first set of wings in the same wild type and hybrid wasps as panel (D). ANOVA analysis reveals no significant asymmetry in legs and wings. (P=0.28 among legs and P=0.65 among wings).

### Alleles causing head defects are recessive whereas alleles governing head shape are codominant

Due to the obligate haplodiploidy, hymenopterans such as *Nasonia*, males are normally hemizygous and interactions among alleles can be assessed in the absence of dominance effects. However, understanding the dominance relationships of alleles is helpful in understanding both the function of the genes involved in generating a phenotype, and the molecular nature of interactions that lead to changes or failure in development.

To study the dominance relationships between the two parental genomes while maintaining male-specific traits, we created diploid males using the previously described method of knocking down the maternal Nv-tra contribution by pRNAi. In the absence of maternal Nv-tra, mated females will produce diploid males (Verhulst *et al.* 2010; Beukeboom *et al.* 2015), *Nv-tra* dsRNA injected *Nv* females were mated to *Ng* males, which resulted in diploid, hybrid male, offspring (Figure 5C). Since these offspring are F1 hybrids where not genetic recombination or assortment has taken place, and there are no sex-based chromosomal differences in these species, they receive an equal contribution genetic material from each parental species (along with the lack of sex chromosomes in this system). Interestingly, we found that for almost all traits, the phenotype for these diploid hybrid males was nearly exactly intermediate between, and significantly different from, both of the parental species (Figure 5A-C, Figure 6, Table S3). Minor deviations from this pattern were at OIO measurements which were closer to those of *N. vitripennis*, while MHW and AIO were more similar to that of *N. giraulti* (Figure 6, Table S3).

**Figure 5.**
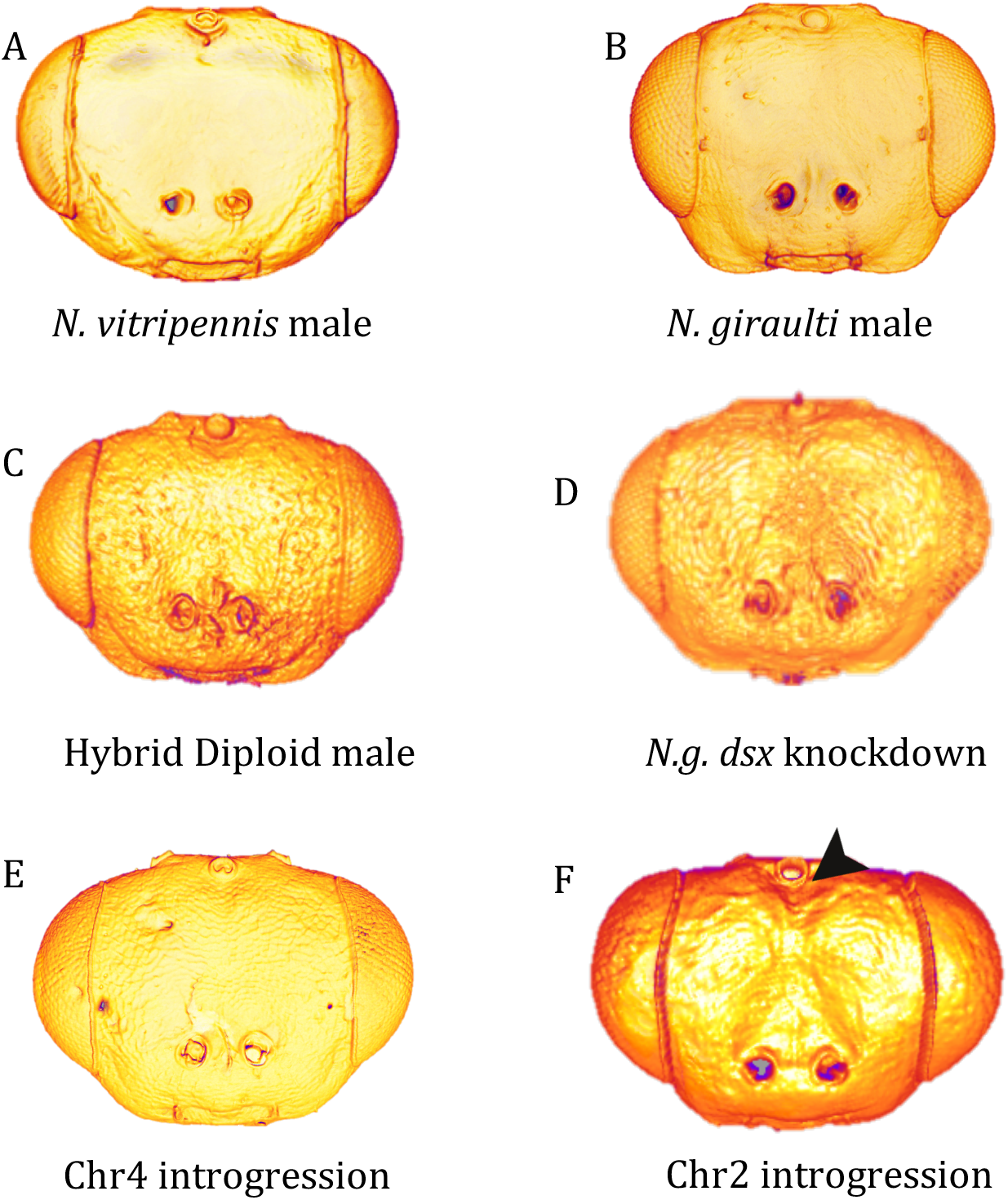
Experimental hybrid head shapes. A) Wild type *N. vitripennis* male B) Wild *type N. giraulti* male, C) Diploid male, D) *N.g. dsx* knockdown, E) Introgression on chromosome 2, F) Introgression on chromosome 4, arrowhead points to midline cleft. Note no other obvious asymmetries or abnormalities.

**Figure 6.**
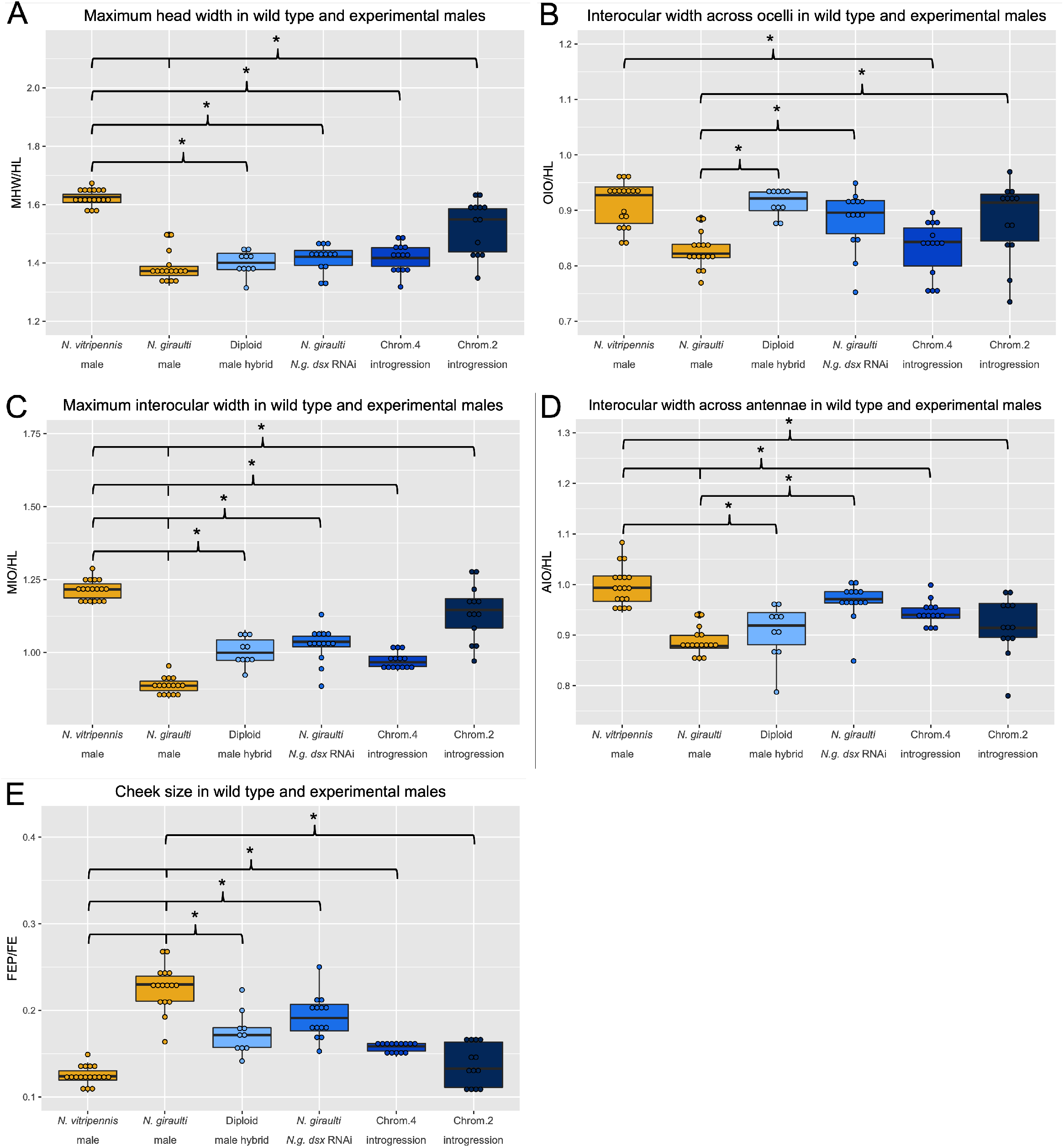
Measurement ratios of RNAi and introgression experiments, presented as box and whisker plots. Each dot represents a single individual, a box represents the inter-quartile range, the center line represents the median value and vertical lines represent upper and lower quartile ranges. A) Maximum head width over head length (MHW/HL), B) Interocular width at ocelli over head length (OIO/HL), C) Maximum interocular widther over head length (MIO/HL), D) Interocular width at antennae over head length (AIO/HL), E) Cheek size (FEP/FE). Wild type *N. vitripennis* and *N. giraulti* males are shown in yellow and experimental lines in varying shades of blue. Each experimental group was compared to both wild type groups. Asterisks indicate P<0.05.

These results indicate that alleles of genes involved in regulating the size and shape of the head are codominant, leading to hybrids with intermediate traits. Additionally, diploid males did not display any of the anomalous phenotypes that occur in haploid hybrids, indicating that it is not the mere presence of an allele from the other species that causes the incompatibility. Rather, it appears that there are species specific alleles that can only function with alleles at other loci that are derived from the same species, and it is the absence of the compatible alleles that leads to defects in the hybrid F2 males between *Nv* and *Ng*. In other words, hybrid head defects involve recessive interaction among loci from the two species.

### Doublesex knockdown in *N. giraulti* males generate a reduced cheek phenotype

Since the divergent head morphology in *N. giraulti* is a specific novelty in the males (the females are barely distinguishable from other *Nasonia* species females), we hypothesized that effectors of the sex determination system may play an important role in generating the divergent features of the *N. giraulti* male head. To test this, we examined the involvement of *doublesex (dsx)* in craniofacial development in *N. giraulti*. Dsx is the main effector gene of the sex determination pathway, and it is known to play specific roles in the evolution of developmental traits that vary between sexes (Hediger *et al.* 2004; Verhulst *et al.* 2010; Tanaka *et al.* 2011; Ito *et al.* 2013), including sex specific differences in wing size between *Nv* and *Ng* (Loehlin *et al.* 2010). We therefore hypothesized that it would play a role in generating the sex specific features of the *N. giraulti* male head. Larval RNAi (Werren *et al.* 2009)was used to knock down *N. giraulti doublesex* (*Ng-dsx*) in male (progeny of virgin females) late-stage larvae before the main period of growth and patterning of the eye and antennal imaginal discs commenced.

Compared to wild type *N. giraulti* males, the divergent *N. giraulti* features of the male were significantly reduced by Ng-dsx knockdown, (Figure 5D, Figure 6B-E, Table S3). MIO and cheek size (FEP/FE) were significantly different from both wild-type *Ng* (p<0.01 and 0.05, respectively) and wt *Nv* (both p<0.01) after Ng-dsx RNAi. OIO and AIO were strongly different from wt *Ng* (p<0.01), but were statistically indistinguishable from *Nv* males, indicating that these features are strongly influenced by *Ng-dsx*. (Figure 6, Table S3). From these results we can conclude that *Ng-dsx* plays an important role in producing the lineages specific male traits in *Ng*. It is possible that female form is the default, and genes that determine male sex also cause the male-specific facial morphology. However, the Ng-dsx RNAi strain was still significantly different from *Ng* females at MHW, MIO and cheek size (Table S4), indicating there was not a complete transformation to the female phenotype, and we can conclude that *dsx* likely works alongside many other genes to generate the male form.

### Introgression of *N. giraulti* dsx non-coding region increases cheek size

The role of *Ng-dsx* in generating the *N. giraulti* male specific structures was further tested by taking advantage of an introgression line containing a portion of the regulatory region of *Ng-dsx* isolated in the background of *N. vitripennis* (Figure 5D). This introgression was originally identified as a region important for the larger size of the *Ng* male wing (Loehlin *et al.* 2010). This relatively small introgression (~40kb) containing only DNA in the non-coding region upstream of the transcription start site of *Ng-dsx* has a strong effect on the shape of the male head in an *N. vitripennis* background. For all five measures examined, the introgression line showed highly statistically significant difference to normal *Nv* male values (p<0.01 for all values, Table S3). Additionally, the introgression line was not statistically significant from normal *Ng* males at MHW and OIO, which is consistent with our hypothesis that *dsx* plays a crucial role in generating the *N. giraulti* specific male head shape features.

Since this introgression line also shows significant differences in shape also from *N. giraulti* (Figure 5E, Figure 6), it is clear that other factors are involved. It is likely that multiple loci contribute significantly to the head shape differences, as seen for the wing size and shape network differences between these two species (Gadau *et al.* 2002). Indeed, complex genetic bases for all of the differing male head shape and size features were predicted in our previous quantitative trait locus analysis (Werren et al 2016). That being said, we cannot exclude that *Ng-dsx* plays a larger role than that detected here. We do not know exactly how *Nv-dsx* expression is being affected in the head, and there may be additional enhancers not included in the introgressed region that are important for additional aspects of dsx expression divergence between the species.

### Introgression of incompatible loci lends insight to abnormal clefting phenotype

As shown above (and previously in Li *et al.* 2005; Loehlin *et al.* 2010; Loehlin and Werren 2012; Hoedjes *et al.* 2014), introgression of genomic regions from one species’ background into another is a powerful method to analyze the genetic basis of evolutionary traits in *Nasonia*. Previous QTL analyses for clefting showed a complex web of genetic interaction among regions on chromosomes 2, 4 and 5 (Werren *et al.* 2016). Briefly, clefting occurs at frequency of ~25% when either or both the regions on Chr 2 and Chr 4 have the *N. giraulti* genotype AND the region on Chr 5 has the *N. vitripennis* genotype. If Chr 5 has the *N. giraulti* genotype, clefting is completely suppressed, unless both the Chr2 and Chr 4 region derives from *N. vitripennis.*Clefting also occurs at about 25% of the time when all three regions derive from *N. vitripennis*, indicating that at least one more locus is involved, or that there is an effect of the general hybrid background on the threshold for clefting.

To simplify analysis of this trait, we examined existing introgression lines with segments of *Ng* DNA introgressed in a *Nv* background. One line, derived from a larger introgression spanning the centromere of chromosome 2 consistently showed facial clefting (See Methods, Figure 5F). Significantly, the females homozygous for this introgression also display the cleft phenotype, unlike F1 hybrid females that never show abnormalities. This shows that the interactions leading to the epistatic phenotype are recessive, since the introgression lines are homozygous and are not seen in the F1 females. The result is consistent with the F2 clefting QTL analysis which predicts that the *Ng* allele in chromosome 2 will induce clefting when combined with the *Nv* alleles at the locus on chromosome 4 or 5 (Werren *et al.* 2016). This result also indicates that the clefting trait is not directly related to the sex specific morphological divergence between the species, and is rather a general defect in head patterning. Finally, this introgression importantly shows that, at least for the locus on chromosome 2, the clefting trait is fully penetrant when incompatible alleles are isolated from any suppressing alleles at other loci. This will simplify identification of the causative allele from *N. giraulti*, and aid in the fine-scale mapping and positional cloning of suppressing alleles at other loci (e.g. on chromosome 5).

## Discussion

In these experiments, we have demonstrated the use of *Nasonia* genus of parasitic wasps to explore the genetic basis of shape. Taking advantage of the significant differences in cranial morphologies among three closely related species and the ability to generate interspecies hybrids, we are able to begin unraveling the network of gene interactions that govern trait formation. Both morphology and genetic incompatibility are result of complex epistatic interactions (Werren *et al.* 2016).

Abnormally asymmetric phenotypes, as seen in the hybrid F2 males here, are known as fluctuating asymmetries, typically caused by developmental instability (Dongen 2006). Developmental instability can result from any number of genetic or environmental factors, commonly observed when hybridizing genomes (Leamy and Klingenberg 2005). However, our system differs from typically cases of fluctuating asymmetry, since we observe stable symmetry among the rest of the body in hybrid males. This indicates generalized developmental instability is not likely to be the cause in this case. This indicates that the head asymmetries we observe in F2 males are not likely to be due to general developmental instability, but rather have a specific genetic basis in the context of head development. The feasibility of dissecting gene interactions governing complex head defects using introgression and recombination mapping has already been shown with our work with the clefting trait, so *Nasonia* is well positioned to make a unique contribution to understanding the genetic and developmental causes of fluctuating asymmetry. Sex identity clearly plays an important role in some aspects of the head shape network, at least in *N. giraulti.* While knockdown of the male-specific spliceoform of *Ng-dsx* does decrease male-specific morphology, a full transformation to the female form may require the function of the female specific transcript of *Ng-dsx.* However our results are also consistent with a complex interplay between sex-specifc genes, and developmental factors shared between the sexes in generating sex specific morphologies. While morphology is strongly influenced by sex, the negative gene interactions that cause developmental defects in F2 hybrid males clearly are not, since our clefting introgression line shows the phenotype in homozygous females as well as haploid males.

QTL analysis is valuable as a starting point for fine-scale mapping of interacting loci that are the genetic basis for observed disrupted phenotypes (Gadau *et al.* 1999). Putative causal regions can be isolated in the other species’ genetic background by introgression for further analysis. Introgression is a very powerful method to understand quantitative traits and gene interactions, whereby a section of one genome is isolated in the background of another through a series of backcrosses, and its localized effects examined. Introgression lines are also powerful starting points for fine scale mapping and positional cloning of causative alleles. The introgression of the clefting locus on chromosome 2 is a good example of the power of the introgression approach. Given the complexity of the interactions that govern the appearance of the cleft in F2 hybrid males, it was somewhat surprising that the introgression of the *N. giraulti* Chr2 locus led to a completely penetrant phenotype in both males and females, behaving basically as a Mendelian recessive allele. Thus it appears that while the genetic architecture preventing clefting in the pure species is complex, each individual allele may have a relatively simple and robust role, rather than each locus having an unpredictable magnitude of effect on the phenotype.

Future analyses will focus on determining whether the other participating alleles predicted by the QTL analyses (Werren *et al.* 2016) also have strong effects in a foreign background, or if there is a mixture of completely, and incompletely, penetrant negative interactions. In particular, a region on Chr5 interacts with the region from Chr2. Based on the QTL analysis (Werren *et al.* 2016), we expect an introgression of the Chr 5 region to completely suppress clefting in combination with the Chr 2 introgession, since clefting occured 0% of the time when these two alleles were present together in F2 males used for the QTL analysis. The expected phenotype of this Chr5 region are less clear, since overall clefting occured 25% of the time when regions on both Chr 2 and Chr4 had the *N. vitripennis* genotype. (Werren *et al.* 2016). This indicates either that there are other loci that suppress clefting induced by the *N. giraulti* Chr5 allele, or that this allele does not promote clefting in a fully penetrant way.

We intend to map these additional interacting loci governing the evolution of morphology by first using Multiplexed Shotgun Genotyping and QTL analysis to identify genomic segments associated with the traits of interest (Andolfatto *et al.* 2011). We can then use marker based introgression and recombination mapping to identify the causative alleles.

In crosses between the closely related flies *Drosophila simulans* and *D. mauritiana* which have divergent head shapes, seemingly coordinated changes in size of the eye field and facial cuticle were found to be due to separable genomic loci (Arif *et al.* 2013). No complex gene interactions or developmental defects (such as clefting or asymmetry) were reported. This may be due to the shorter divergence time between the *Drosophila* species (~250,000 years (ref: Genome Res. 2012. 22: 1499-1511)) than between *N. vitripennis* and *N. giraulti* (~1 million years). Our results are consistent with the appearance of negative epistatic interactions between isolated species being correlated with increased time since divergence, since we do not observe these effects in hybrids of closely related species *N. longicornis* and *N. giraulti.*Future analysis of the genetic architecture of the morphological difference between *N. longicornis* and *N. giraulti* also have more simple genetic bases, like those observed between *D. simulans* and *D. mauritiana*, or whether epistasis plays an important role already in more recently diversified species of *Nasonia.*

In conclusion, our results have demonstrated important roles of sex, ploidy, and divergence time in the evolution of novel morphologies and developmental defects in hybrids. In addition, our introgression of an allele from one species that causes a severe developmental defect in the genomic background of its close relative is an important step in simplifying an understanding of the still daunting task of characterizing gene interactions involved in head development and developmental abnormalities. The powerful genetic tools available in *Nasonia* wasp combined with the rich, complex genetic architectures of the head shape differences and developmental defects, will make these parasitoids excellent models for charting the connections between genomic and phenotypic variation.

